# A rapid and sensitive multiplex, whole mount RNA fluorescence *in situ* hybridization and immunohistochemistry protocol

**DOI:** 10.1101/2023.03.09.531900

**Authors:** Tian Huang, Bruno Guillotin, Ramin Rahni, Ken Birnbaum, Doris Wagner

**Affiliations:** University of Pennsylvania, Department of Biology, Philadelphia, PA 19104, USA; Department of Biology, Center for Genomics and Systems Biology, New York University, New York, NY 10003, USA

**Keywords:** RNA-FISH, hybridization chain reaction, whole mount, immunohistochemistry, fluorescent protein

## Abstract

In the past few years, there has been an explosion in single-cell transcriptomics datasets, yet *in vivo* confirmation of these datasets is hampered in plants due to lack of robust validation methods. Likewise, modeling of plant development is hampered by paucity of spatial gene expression data. RNA fluorescence *in situ* hybridization (FISH) enables investigation of gene expression in the context of tissue type. Despite development of FISH methods for plants, easy and reliable whole mount FISH protocols have not yet been reported. We adapt a 3-day whole mount RNA-FISH method for plant species based on a combination of prior protocols that employs hybridization chain reaction (HCR), which amplifies the probe signal in an antibody-free manner. Our whole mount HCR RNA-FISH method shows expected spatial signals with low background for gene transcripts with known spatial expression patterns in Arabidopsis inflorescences and monocot roots. It allows simultaneous detection of three transcripts in 3D. We also show that HCR RNA-FISH can be combined with endogenous fluorescent protein detection and with our improved immunohistochemistry (IHC) protocol. The whole mount HCR RNA-FISH and IHC methods allow easy investigation of 3D spatial gene expression patterns in entire plant tissues.

## Introduction

Profiling spatiotemporal gene expression patterns is critical for studying developmental biology. RNA fluorescence *in situ* hybridization (FISH) allows detection of spatial gene expression at different developmental stages of an organism. Several types of FISH methods have been established for plants (Rozier et al., 2014; Duncan et al., 2016; Huang et al., 2020; Solanki et al., 2020; Yang et al., 2020). Traditionally, oligonucleotide probes targeting specific transcripts are labeled by epitopes. After hybridizing to the target RNA, the probes are visualized by antibody-based methods (Rozier et al., 2014; Yang et al., 2020). Alternatively, for single molecule FISH (smFISH), short oligonucleotide probes targeting RNA can be conjugated with fluorescent dyes, and the probes visualized directly after hybridization (Duncan et al., 2016; Huang et al., 2020). Tens of short smFISH probes are designed for one RNA, which allows sensitive detection of a single RNA molecule. Recently, a FISH method with branched DNA (bDNA) amplification was also described in plants (Solanki et al., 2020). Despite these advances, most existing RNA FISH methods still require sectioning of the tissue, and the samples are processed and visualized on slides, which is laborious and only provides spatial information in two dimensions (Duncan et al., 2016; Huang et al., 2020; Solanki et al., 2020; Yang et al., 2020). Rapid and robust methods are needed in plants to corroborate or identify the tissue of origin for single-cell clusters but also generally speed up the workflow of *in situ* hybridization.

RNA-FISH based on hybridization chain reaction (HCR) has been designed, tested, and optimized in animal species (Choi et al., 2010; Choi et al., 2014; Choi et al., 2016; Choi et al., 2018; Trivedi et al., 2018). HCR enables antibody-free FISH signal amplification via the self-assembly of small oligonucleotides (Dirks and Pierce, 2004). A recently improved HCR RNA-FISH method (HCR RNA-FISH v3) was reported to have higher sensitivity and robustness with background suppression in all steps (Choi et al., 2018). The ease of multiplexing different HCR probe sets also allows simultaneous detection of multiple RNA species. Furthermore, since no protein is involved in this method, it alleviates possible problems with protein penetration in thick tissues, making whole mount FISH much more feasible.

In this paper, we describe a simple 3-day whole mount RNA-FISH protocol for *Arabidopsis thaliana* (Arabidopsis), *Zea mays* (maize), and *Sorghum bicolor* (Sorghum) using HCR. This protocol allows high throughput processing of samples in Eppendorf tubes with limited handling, low hybridization temperature, and probe signal that persists for several days after processing if samples are stored at 4°C. We show that HCR RNA-FISH can detect known gene expression in whole mount plant tissue—even for genes that are expressed in deep tissue layers—and that we can monitor at least two or three genes simultaneously in maize/Sorghum and Arabidopsis, respectively. Additionally, this protocol allows the preservation and detection of expressed fluorescent proteins such as GFP alongside FISH probe signal. Finally, we establish an improved protocol for combined FISH and immunohistochemistry (IHC) that can detect RNA and protein in the same sample. This greatly facilitates the study of mobile proteins or of transcription factors and their targets.

## Results

### Development of a whole mount FISH protocol

We combined and optimized two previously described protocols (Rozier et al., 2014; Choi et al., 2018) for whole mount RNA-FISH. To achieve better probe penetration, the cuticle, cell membrane, and cell wall of fixed plant samples are permeabilized through alcohol treatment and cell wall enzyme digestion (Rozier et al., 2014; Young et al., 2020) (Fig. 1a). HCR RNA-FISH is performed on fixed, permeabilized plant samples according to previously described methods in animal species (Choi et al., 2018) (Fig. 1a). Briefly, probe sets contain multiple hybridization probe pairs that bind different sites on the RNA target. Each probe pairs consist of two small 25 nucleotide single strand DNA probes hybridizing on adjacent sequences of the target mRNA, and each probe contains half of a small DNA initiator sequence. Only when both probes hybridize next to each other can the split-initiators form an intact initiator (Fig. 1b). The initiator triggers the self-assembly of hairpin amplifiers which are tagged by fluorescent dyes, leading to an amplification of fluorescent signal (Fig. 1c). By multiplexing different initiator/amplifier sequences (e.g. B1, B2, B3…) and different fluorescent dyes, simultaneous detection of multiple RNA targets in the same sample can be easily achieved.

**Figure 1.**
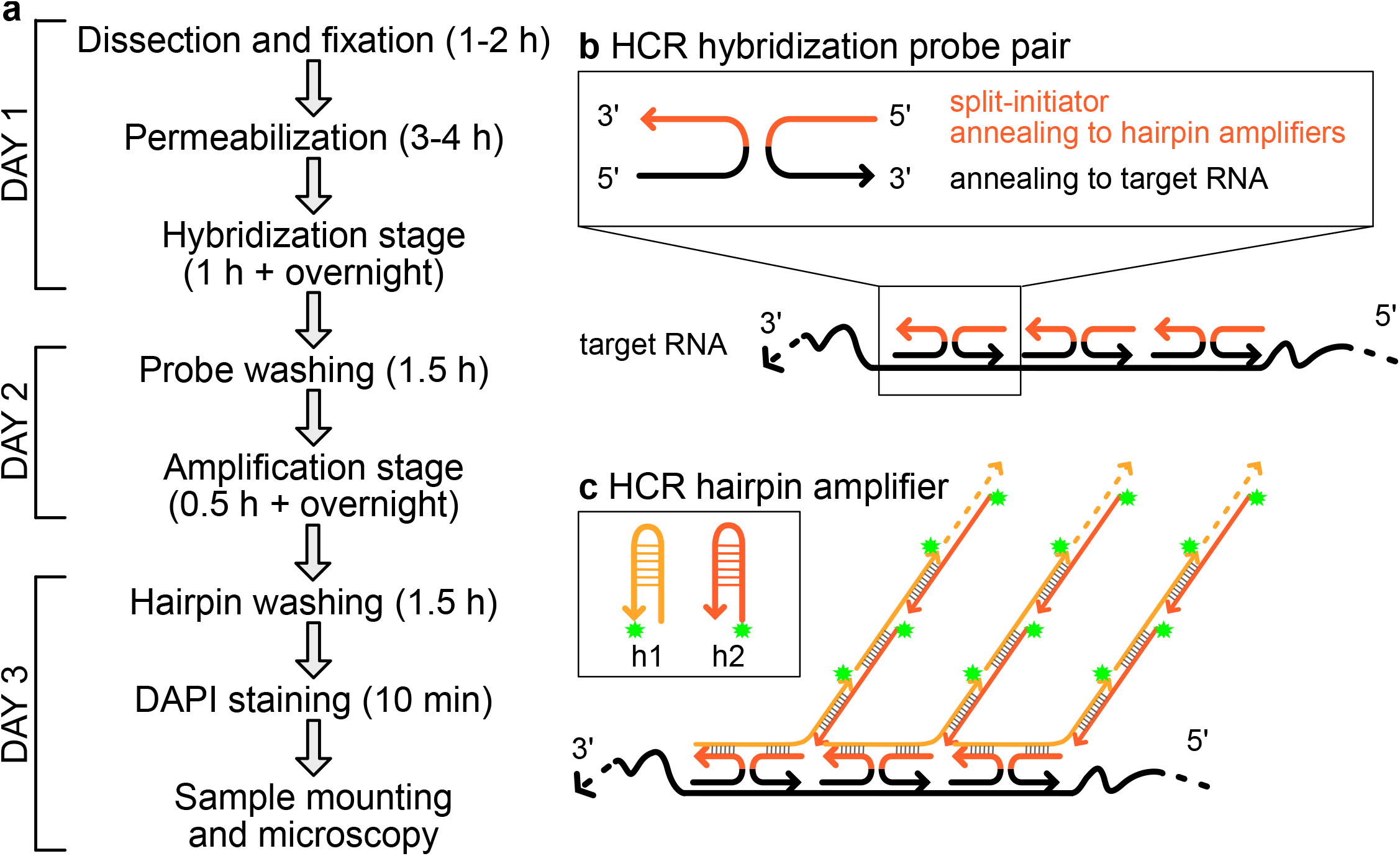
Schematic of plant wholemount HCR RNA-FISH workflow and mechanism. a. Timeline of 3-day plant wholemount HCR RNA-FISH protocol. b. In the hybridization stage, HCR hybridization probes anneal to the target RNA, and adjacent probes form an initiator which allows the initiation of the HCR amplification. c. In the amplification stage, HCR hairpins (h1 and h2) stay self-annealing in the absence of an initiator. When an initiator is present, hairpin h1 and hairpin h2 hybridize to each other and initiate the self-assembly (hybridization chain reaction). Hairpins are tagged by fluorescent dyes (green star), and the self-assembly leads to an amplification of fluorescent signals.

To test whether HCR RNA-FISH can detect gene transcripts with known spatial expression pattern, we first chose to examine the expression of the stem cell regulators *CLAVATA3* (*CLV3*) and *WUSCHEL* (*WUS*) in Arabidopsis inflorescences. *CLV3* is expressed in the stem cell niche in the center of the shoot apex, while *WUS* is expressed in the organizing center region below *CLV3* (Mayer et al., 1998; Fletcher et al., 1999). HCR RNA-FISH allowed simultaneous detection of both *WUS* and *CLV3* in a single inflorescence when viewed from above (Fig. 2a). Double labeling showed that *WUS* expression starts to appear in stage 1 flower primordia (Fig. 2a, arrowhead), while *CLV3* expression appears later in stage 2 flower primordia (Fig. 2a, arrow) (flower stages are determined according to (Smyth et al., 1990)). This agrees with previous observations of *WUS* and *CLV3* temporal expression patterns in flower primordia (Mayer et al., 1998; Fletcher et al., 1999). Optical longitudinal sections and 3D projection revealed that the *WUS* domain was below the *CLV3* domain, as previously reported (Gruel et al., 2016) (Fig. 2a,c). As a negative control, we employed probes targeting mScarletI and mEGFP and observed a low level of autofluorescence with minimal non-specific binding and amplification (Fig. 2b). Non-specific uniform background was slightly stronger for Alexa Fluor 488 (green) than for Alexa Fluor 546 (red). We also detected *WUS* and *CLV3* expression in young Arabidopsis shoot apical meristems during the floral transition using a “half mount” protocol (Fig. 2d). 11-day-old plants were sectioned longitudinally by razor blade through the center of the plant before RNA-FISH. Similar to the inflorescence meristem, *CLV3* signal is present above the *WUS* domain, as expected.

**Figure 2.**
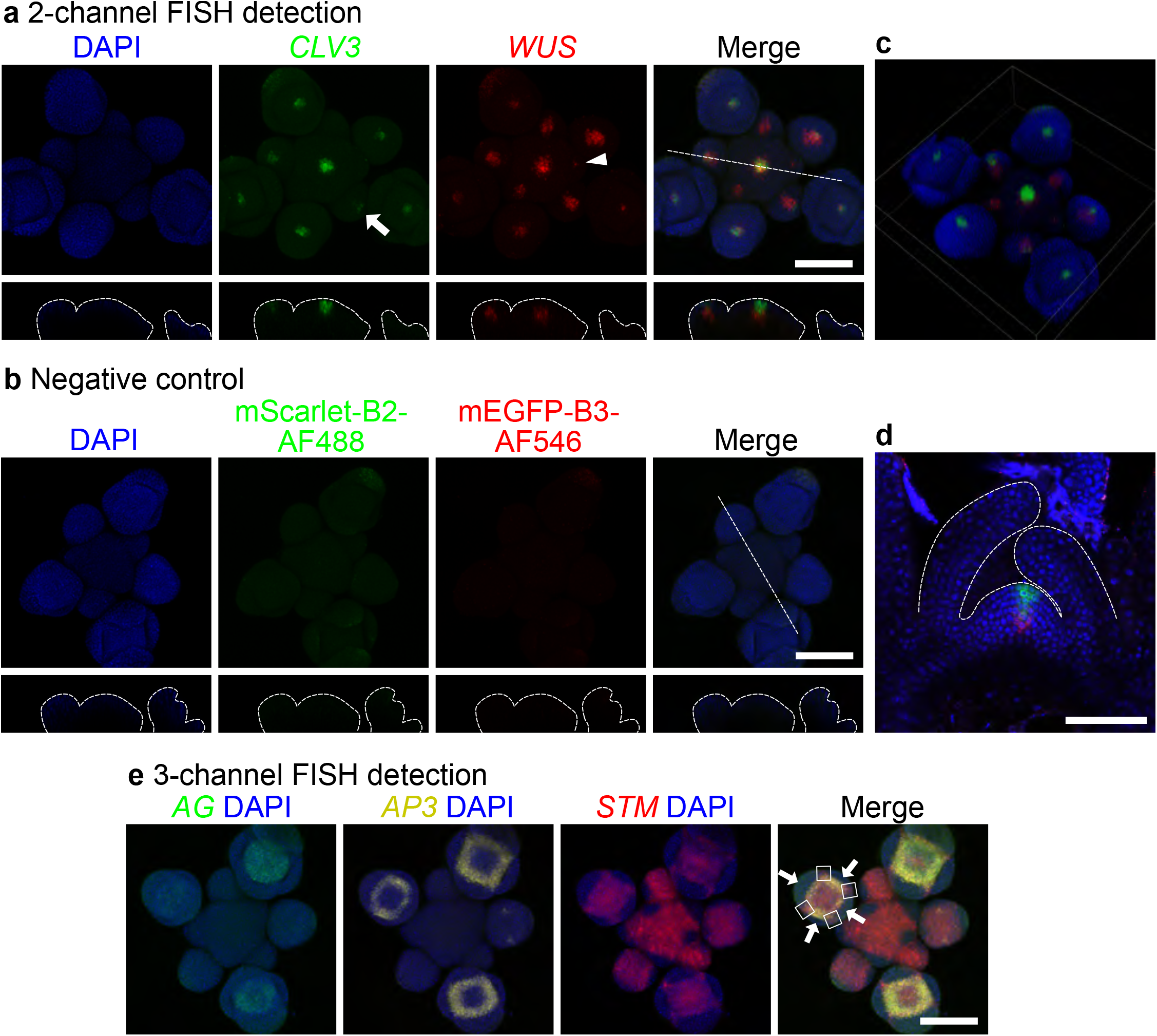
Multiplexed HCR RNA-FISH in wildtype Arabidopsis inflorescence. a. 2-channel FISH for *CLV3* (green) and *WUS* (red) in inflorescence (top-view with maximum intensity projection). HCR hairpin amplifiers B2-AlexaFlour488 (B2-AF488) and B3-AlexaFlour546 (B3-AF546) were used in the amplification stage. The arrow and arrowhead indicate the earliest flower primordia that express *CLV3* and *WUS* respectively. The orthogonal views across the dash line were shown in the bottom. b. Test for background. All FISH steps were same as those in Fig. 2a except that mScarletI-B2 and mEGFP-B3 were used in the hybridization step as negative controls. The orthogonal views across the dashed line were shown in the bottom. c. 3D projection of the sample in panel a. d. FISH for *CLV3* (green) and *WUS* (red) in 11-day-old shoot apical meristem (side-view). e. 3-channel FISH for *AG* (green), *AP3* (yellow), and *STM* (red) in inflorescence (top-view with maximum intensity projection). White arrows indicate the lateral and medial sepal primordia. White squares represent the *STM* expression domain between adjacent sepal primordia. Nuclei were stained by DAPI (blue). Scale bar = 100 μm.

Next, we tested the capability of HCR RNA-FISH to detect 3 transcripts simultaneously in the same inflorescence. *APETALA* 3 (*AP3*), *AGAMOUS* (*AG*), and *SHOOT MERISTEMLESS* (*STM*) were simultaneously probed and detected in the same inflorescence with previously reported spatial expression pattern (Yanofsky et al., 1990; Jack et al., 1992; Long et al., 1996; Long and Barton, 2000; Krizek and Fletcher, 2005) (Fig. 2e). *AP3* expression occurred one primordium prior to that of *AG. AG* expression partially overlapped with that of *AP3*, as expected since *AP3* and *AG* together specify stamen identity (Goto et al., 2001). *STM* expression in the meristem was excluded from incipient and very young (<stage 1) flower primordia, as expected (Long et al., 1996; Long and Barton, 2000). *STM* was expressed in older flower primordia, but was excluded from the lateral and medial sepal primordia (Fig. 2e, white arrows), as previously observed (Long and Barton, 2000). From stage 3 onward, *STM* expression in flower primordia was entirely contained within the regions demarcated by *AP3* and *AG*, with the exception of the boundaries between the sepal primordia (white squares, see also Long 2000 (Long and Barton, 2000)).

In conclusion, the results of HCR RNA-FISH show expected spatiotemporal gene expression pattern with low background in Arabidopsis inflorescences. Also, HCR RNA-FISH allows simultaneous detection of 2 or 3 different gene transcripts in the same sample.

### Simultaneous FISH and detection of endogenous fluorescent reporters

To test whether HCR RNA-FISH can be used together with fluorescent reporters, we monitored transgene expression in null mutants for the *TERMINAL FLOWER 1* (*TFL1*) gene rescued by a translational protein fusion to the EGFP fluorescent protein (gTFL1-GFP *tfl1-1* (Zhu et al., 2020)) as it is known that TFL1 protein moves beyond its site of transcription (Conti and Bradley, 2007; Goretti et al., 2020). For FISH we used probes targeting *EGFP* mRNA and—to avoid possible bleed-through between channels—we chose probes with a fluorescent dye (Alexa Fluor 546) whose excitation and emission spectra do not overlap with EGFP fluorescence. *EGFP* mRNA was detected in the center of the meristem in gTFL1-GFP *tfl1-1*, while little signal was observed in wild type plants using the same probes, as expected (Fig. 3b). This absence of off target binding highlights the high specificity of the signal detected using HCR FISH. The pattern of *EGFP* transcript in gTFL1-GFP *tfl1-1* resembled that of the endogenous *TFL1* transcript in the wild type, which is restricted to the center of the inflorescence meristem (Bradley et al., 1997). Interestingly, despite methanol and ethanol dehydration in the FISH protocol, we were still able to detect EGFP fluorescence, albeit at lower intensity than in tissues not subjected to FISH. Nevertheless, gTFL1-GFP fluorescence was detected in the previously reported protein accumulation domain (Zhu et al., 2020) simultaneously alongside *EGFP* mRNA. Simultaneous detection of protein fluorescence and RNA allows direct comparison of the protein and RNA expression domains of mobile proteins—like TFL1—which are common in plants (Barton, 2001; Conti and Bradley, 2007; Goretti et al., 2020). It also allows simultaneous visualization of other reporters (for hormones, subcellular compartments, etc.) alongside transcripts. Conversely, to avoid potential overlap between the excitation and emission spectra of the fluorescent protein and the FISH probe, the fluorescent protein can be easily removed by treating with proteinase K (Fig. 3).

**Figure 3.**
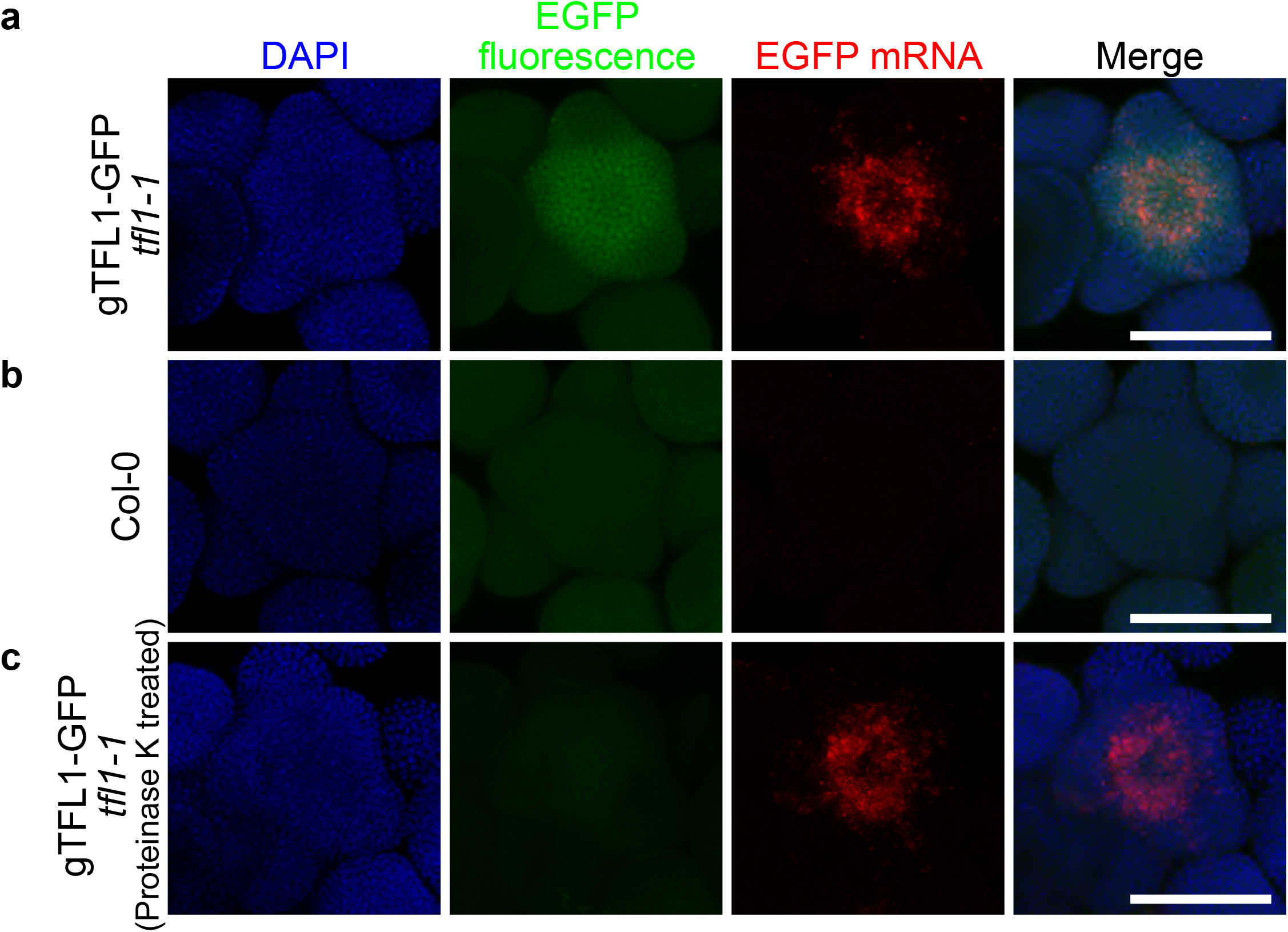
HCR RNA-FISH in the inflorescence of Arabidopsis transgenic reporter. a-b. EGFP fluorescence (green) and RNA-FISH for *EGFP* transcripts (red) were detected in gTFL1-GFP *tfl1-1* (a), Col-0 wild type (b) (top-view with maximum intensity projection). The detection in Col-0 wild type served as negative control. c. EGFP fluorescence (green) and FISH for EGFP transcripts (red) in gTFL1-GFP *tfl1-1* with proteinase K treatment (top-view with maximum intensity projection). Nuclei were stained by DAPI (blue). Scale bar = 100 μm.

### Combined FISH and IHC

Combining RNA-FISH with fluorescence detection allows co-detection of RNA and proteins fused to fluorescent proteins. However, it is not compatible with proteins labelled by small epitope tags (e.g. HA, Myc) and endogenous proteins without tags. Whole mount immunohistochemistry (IHC) methods have been described for Arabidopsis inflorescence (Pasternak et al., 2015; Tran et al., 2021). IHC allows detection of proteins whenever an antibody against the target protein or the epitope tag is available. To test whether HCR RNA-FISH can be combined with IHC, we attempted to simultaneously detect *EGFP* mRNA and protein in gTFL1-GFP *tfl1-1* using *EGFP* RNA-FISH and anti-GFP IHC (Fig. 4). A standard HCR RNA-FISH was performed to detect *EGFP* mRNA. Next, extra cell wall digestion and post-fixation were applied to the samples to achieve higher permeability for antibodies in the IHC. Blocking, primary antibody incubation, and secondary antibody incubation were then performed in a similar way described in published IHC methods. We chose Alexa Fluor 514 for RNA-FISH and Alexa Fluor 546 for secondary antibody in IHC so that the spectra of both fluorescent dyes do not overlap with each other and do not overlap with EGFP fluorescence. As shown in Fig. 4, EGFP protein was detected by anti-GFP IHC (red) in the meristem of gTFL1-GFP *tfl1-1*, and only a weak background was detected in the Col-0 wild type negative control. We noticed EGFP fluorescence (green) was also partially maintained, and the pattern of IHC signal (red) resembled that of the EGFP fluorescence (green). At the same time, *EGFP* mRNA was detected in gTFL1-GFP *tfl1-1* by HCR RNA-FISH (Fig.4, yellow). In the Col-0 wild type negative control, EGFP fluorescence (green) and RNA (yellow) were not detectable, as expected. These results suggested that multiplexed IHC and HCR RNA-FISH had good preservation of FISH signal and showed expected IHC signal pattern. Although we chose EGFP as the target for IHC detection, all kinds of epitope tags should be compatible with this method as long as the chosen primary antibody shows little non-specific binding (low background). When antibodies against endogenous proteins are available, it is possible to simultaneously probe the endogenous proteins with mRNA targets in non-transgenic plants. In summary, we successfully combined the HCR RNA-FISH protocol with immunohistochemistry which allows simultaneous detection of RNA and protein in whole mount tissue.

**Figure 4.**
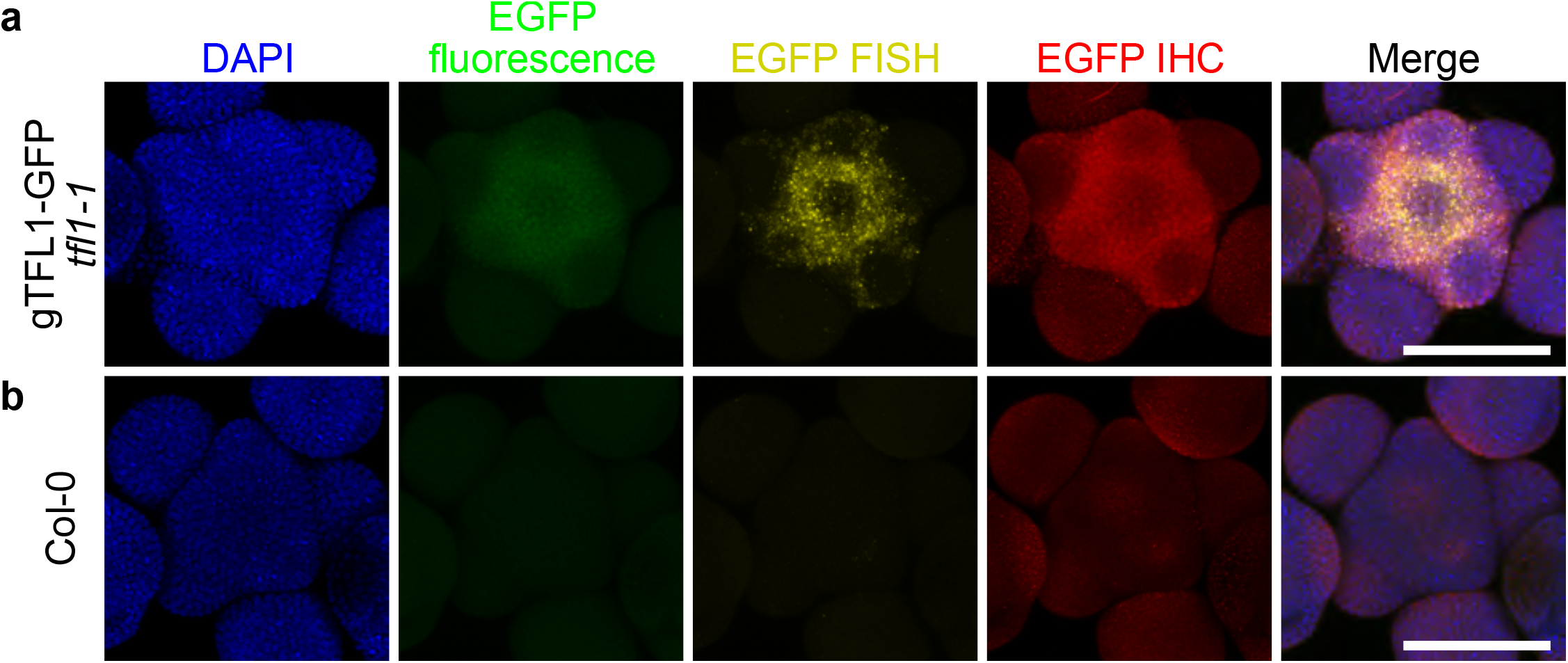
Combined FISH and IHC in Arabidopsis inflorescence. EGFP fluorescence (green), RNA-FISH for *EGFP* transcripts (yellow), and anti-GFP IHC were detected in gTFL1-GFP *tfl1-1* (a), Col-0 wild type (b) (top-view with maximum intensity projection). The detection in Col-0 wild type served as a negative control. Nuclei were stained by DAPI (blue). Scale bar = 100 μm.

### “Half Mount” FISH in monocot roots

Traditional *in situ* hybridization in monocot roots (Matsuyama et al., 1999) has involved sectioning, which is challenging and time consuming, and whole mount protocols are not currently available for roots. Here, we adapted the HCR protocol on 3D root tissue in *Zea mays* (maize) and *Sorghum bicolor* (Sorghum). Maize and Sorghum seedlings were grown on germination paper, and a few drops of fixative was applied directly to root tips with a pipette, just prior to hand-sectioning along the longitudinal or transverse axis using a microscalpel. Then the tip is excised and directly transferred into fixative solution. While the fixative solution as well as the processing steps prior to HCR differed for monocot samples (see Materials and Methods), the HCR RNA-FISH steps closely follow that described for Arabidopsis above. The resulting “half mount” protocol allowed for clear visualization of HCR probe signal and owing to maize and Sorghum roots’ cell wall autofluorescence, no DAPI counterstain was necessary.

Probes can be visualized with several Alexa fluorophores (Alexa Fluor 488, 514, 546, 594 and 647). Thus, we first tested tissue autofluorescence at the excitation wavelengths for each of the fluorophores, determining that excitation of Alexa Fluor 647 and 488 generated the lowest background fluorescence. Monocots cell walls were visualized using the strong autofluorescence of the tissue under 405nm laser excitation.

To test if HCR RNA-FISH can detect maize transcripts with known spatial expression patterns, we examined ZmGRP4 which has reported expression in the lateral root cap and epidermis (Matsuyama et al., 1999). We reliably detected strong signal from these tissues, whereas negative control probes targeting GFP showed little to no background signal (Fig. 5a-b). In addition, a probe against SCR1h had an expression pattern matching published *in situ* hybridization data (Fig. 5c) (Ortiz-Ramirez et al., 2021).

**Figure 5.**
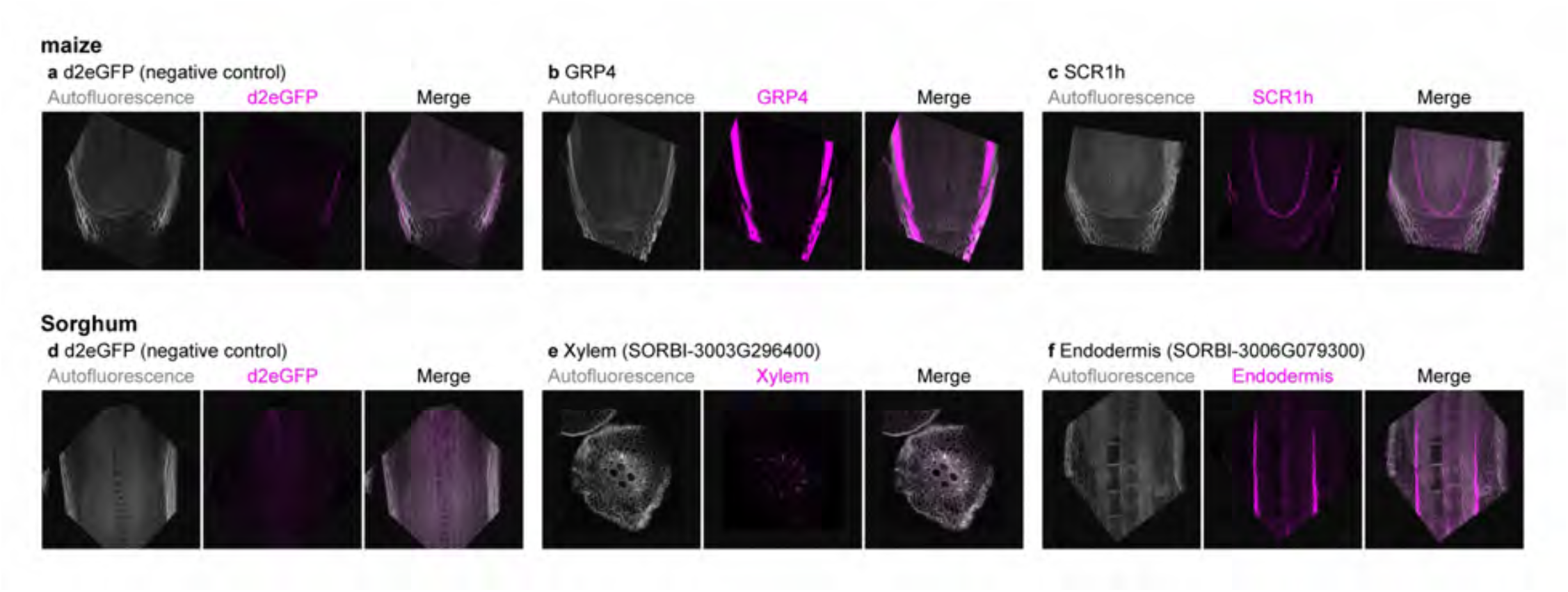
HCR RNA-FISH in monocot roots. HCR RNA-FISH signal in magenta, and cell wall autofluorescence in gray, in either maize (a-c) or sorghum (d-f) roots. a,d FISH background fluorescence was assessed using RNA-FISH for *eGFP* transcripts, not expressed in the roots. b,c longitudinal hand sectioning of maize root tips, revealing the signal for FISH against GRP4 or SCR1h maize genes, specifically expressed in lateral root cap/epidermis (b) or endodermis (c). (e) transversal and longitudinal (f) hand section of sorghum root, RNA-FISH for SORBI-3003G296400 specifically expressed in xylem (e) or for SORBI-3003G079300 expressed in endodermis (f).

Deploying the same protocol in Sorghum, we were able to visualize Xylem and Endodermis markers predicted from single cell (Birnbaum et al., 2022), with negative control probes showing little to no background signal (Fig. 5d-e).

Thus, this protocol allows for a quick and easy imaging of RNA probes in maize and Sorghum root tissue without the need for microtome sectioning. The protocol, including the half mount procedure, greatly speeds up *in situ* hybridization experiments and allows for a sensitive readout of spatial gene expression, even in normally optically inaccessible thick tissues.

## Discussion

With the rapid development of single-cell transcriptomics techniques in plants (Seyfferth et al., 2021; Shaw et al., 2021), our knowledge of tissue specific expression of known regulators has increased dramatically, and an increasing number of unknown genes have been identified that are expressed in specific cell types. Orthogonal approaches are needed to validate these findings and derive biological meaning. This increases the need for testing gene expression patterns by *in situ* hybridization-based methods. Thus far, most approaches used rely on thin sections (Brewer et al., 2006; Duncan et al., 2016; Huang et al., 2020; Solanki et al., 2020; Yang et al., 2020), which preclude 3D visualization of transcript accumulation and greatly add to the time it takes to perform localization assays. Here we provide a robust, versatile, and facile wholemount RNA FISH method based on hybridization chain reaction, which provides a fast and reliable readout for investigating gene expression pattern in complex plant tissues. The method is rapid and allows for a highly sensitive and specific readout of transcript localization. In addition, unlike existing wholemount in situ hybridization methods (Hejátko et al., 2006; Rozier et al., 2014), it allows simultaneous detection of multiple transcripts. We demonstrate that HCR RNA-FISH detects gene expression with precise spatial pattern and low background. Multiplexed detection of different gene transcripts is straightforward due to the nature of HCR technology. Also, the partial persistence of fluorescent protein signal and combined FISH and IHC allows co-detection of the transcript and protein of a gene as we show for the mobile protein TFL1.

To obtain high quality images for HCR RNA-FISH, the most critical steps are sample dissection and mounting. Although HCR RNA-FISH allows detection in deeper regions of tissue, the imaging depth is still restricted to about 100 μm due to light absorption and scattering (Jonkman et al., 2020). Thus, sample dissection is critical, although we note that all dissections in both Arabidopsis and monocot species shown here were performed by hand under a stereoscope scope. In addition, after completing the HCR RNA-FISH procedures, smaller samples tend to be very fragile. Thus, sample handling during the mounting step is critical to avoid damage to the tissue of interest. We also note that sample movement during image acquisition could happen when imaging Arabidopsis inflorescence not well inserted to agarose gel plates. The 3-day HCR RNA-FISH method we describe here will advance elucidation of spatial gene expression pattern in plant species.

## Methods

### Hybridization probes and amplification hairpins

Hybridization probes with split-initiators (e.g. B1, B2, B3) targeting specific transcripts were designed and manufactured by Molecular Instruments. Amplification hairpins with certain amplifier sequences (e.g. B1, B2, B3) and fluorescent dyes were purchased from Molecular Instruments. Probes can be designed to be compatible with Alexa Fluor 488, 514, 546, 594 or 647. All hybridization probes and amplification hairpins used in this study were listed in Table 1. mEGFP probes were used to detect EGFP transcripts since mEGFP and EGFP only have 1 nt difference in sequence.

**Table. 1.** Hybridization probes

| Gene | Initiator/Amplifier-Fluorescent dye | Lot Number |
| --- | --- | --- |
| *WUS* (AT2G17950) | B3 Alexa Fluor 546 | PRM274 |
| *CLV3* (AT2G27250) | B2 Alexa Fluor 488 | PRM273 |
| AG (AT4G18960) | B2 Alexa Fluor 488 | PRM277 |
| AP3 (AT3G54340) | B3 Alexa Fluor 514 | PRM276 |
| STM (AT1G62360) | B1 Alexa Fluor 546 | PRM275 |
| mEGFP | B3 Alexa Fluor 546 or 514 | PRK551 |
| mScarlet-I | B2 Alexa Fluor 488 | PRN494 |
| d2eGFP | B1 Alexa Fluor 647 | PRA221 |
| ZmGRP4<br>(Zm00001d004728/GRMZM2G025205) | B1 Alexa Fluor 647 | PRK923 |
| ZmSCR1h(Zm00001d052380) | B1 Alexa Fluor 647 | PRK918 |
| SORBI-3003G296400 (xylem) | B1 Alexa Fluor 647 | PRK925 |
| SORBI-3006G079300 (endodermis) | B1 Alexa Fluor 647 | PRM013 |

### Reagents

Fixative solutions: (Arabidospis) 4% paraformaldehyde (PFA) (SIGMA, P6148) in phosphate buffered saline (PBS); (Monocot) FAA: 4% formaldehyde, 5% glacial acetic acid, 50% ethanol in RNAse free water.

50% Histo-Clear II / 50% ethanol: Histo-Clear II (Electron Microscopy Sciences, 64111-01) and ethanol were mixed at 1:1 ratio.

DPBST: fresh-made DPBS with 0.1% Tween-20 (Bio-Rad, 170-6531). DPBS was prepared from 10x DPBS without calcium and magnesium (gibco, 14200-075).

5x SSCT: fresh-made 5x SSC buffer with 0.1% Tween-20 (Bio-Rad, 170-6531). 5x SSC buffer was prepared from 20x SSC buffer (CORNING, 46-020-CM).

Cell wall digestion enzyme mix A: The cell wall enzyme mix A formula was adapted from previous publication (Rozier et al., 2014). First, to prepare 6x cell wall digestion enzyme mix A stock, 50 mg Macerozyme R-10 (RPI, M22010), 50 mg Cellulose RS (RPI, C32400), 25 mg Pectolyase (SIGMA, P3026), and 1 mL Pectinase (SIGMA, P4716) were dissolved in 10 mL pure water. The stock was filtered through 0.22 μm syringe filter and stored at -20 °C freezer. To prepare 1x cell wall digestion enzyme mix A, 6x stock was diluted in DPBST.

Cell wall digestion enzyme mix B (for IHC): The cell wall enzyme mix B formula was adapted from previous IHC protocols (Pasternak et al., 2015; Tran et al., 2021). 2x cell wall digestion enzyme mix B stock (0.4% Driselase (SIGMA, D8037) and 0.3% Macerozyme R-10 (RPI, M22010) in PBS) was prepared and stored at -20 °C freezer. 2x stock was diluted in DPBST to prepare 1x cell wall digestion enzyme mix B.

Proteinase solution: 4 μL proteinase K (NEB, P8107S) was added into 1 mL 0.1 M Tris-HCl 0.05 M EDTA (pH 8.0) to prepare the proteinase solution. The buffer (0.1 M Tris-HCl 0.05 M EDTA pH 8.0) was prepared fresh using 1 M Tris-HCl (pH 8.0) (Invitrogen, 15568-025) and 0.5 M EDTA (pH 8.0) (Invitrogen, 15575-020).

DAPI staining solution: 1 μg/mL DAPI (SIGMA, D9542) in DPBS. Blocking buffer: 2% bovine serum albumin (SIGMA, A3059) in DPBST.

Primary antibody solution: diluted primary antibody in blocking buffer. For EGFP IHC, rabbit anti-GFP antibody (abcam, ab290) was diluted at 1:2000 in blocking buffer.

Secondary antibody solution: diluted secondary antibody in blocking buffer. For EGFP IHC, Alexa Fluor 546 goat anti-Rabbit IgG antibody (Invitrogen, A11035) was diluted at 1:200 in blocking buffer.

### Plant growth conditions

Arabidopsis plants were grown in soil at 22 °C under long-day photoperiod (16 h light/8 h dark with light intensity of 120 μmol/m^2^s). gTFL1-GFP *tfl1-1* was described in previous publications [26]. All Arabidopsis plants in this study are Columbia-0 ecotype.

Maize and Sorghum seedlings were surface sterilized for 20mins with 6% active chloride, washed with sterile water then grown on germination paper (Anchor Paper&Cie., 38# regular) in tap water (28°C/24°C, 16 h light/8 h dark with light intensity of 420 μmol/m^2^s) for 7 days.

### Sample dissection and fixation

#### Arabidopsis

Shoot apices were collected shortly after bolting. All flowers covering the inflorescence meristem were dissected and removed by forceps or a needle. It is critical to expose the tissue of interest as much as possible, otherwise confocal laser scanning microscope will not be able to capture signals from tissue of interest. The fixative solution (4% PFA in PBS) was prepared in 1.5 mL centrifuge tube in a fume hood. Samples can be fixed with FAA as well, but FAA might reduce the fluorescent protein signal when co-detecting RNA and fluorescent protein signals. Samples were collected in the fixative immediately after dissection. After vacuum infiltration, samples were fixed for another 30 min at room temperature in the fixative. Fixative was removed by washing once in DPBS for 10 min.

#### Monocots

The fixative solution (FAA) was prepared in 5 mL tube in a fume hood. Just prior to fixation a small volume of fixative FAA was applied directly to the roots using a pipette. Using a microscalpel, roughly 1 cm longitudinal cuts were made in the root tissue, before excising 1.5-2 cm of the root and transferring to fixative solution. Transverse sections, by contrast, were performed just prior to imaging. In fume hood, apply a gentle vacuum until roots float up. Release vacuum, agitate tube, and apply vacuum again. Repeat several times until roots no longer float up (may take up to an hour). Make sure samples are in FAA at room temperature for at least 1 hour. Samples can also be stored in FAA overnight at 4°C.

### Sample permeabilization Arabidopsis

Cuticle and cell membrane can be permeabilized by series of methanol and ethanol incubation (Young et al., 2020). Steps for sample permeabilization was modified from previous FISH protocols (Rozier et al., 2014).

1. Fixed and washed samples were directly dehydrated in methanol twice for 10 min. **PAUSE POINT: samples can be stored in methanol at -20°C for days before continuing**.
2. Then samples were incubated in ethanol twice for 10 min.
3. Samples were cleared and permeabilized in 50% Histo-Clear II / 50% ethanol for 30 min.
4. Samples were then washed in ethanol twice for 10 min and in methanol three times for 5 min.
5. Samples were rehydrated by sequential washing in 75%, 50%, 25%, 0% methanol in DPBST (5 mins each)
6. Partial cell wall digestion was then performed by previously described enzyme mix (Rozier et al., 2014). Samples were incubated for 3 min at room temperature in 1x cell wall digestion enzyme mix A in DPBST. The incubation time for cell wall digestion needs to be adjusted for different tissues, and excessive cell wall digestion often lead to damaged sample structures and diminished FISH signals according to our experience. (Cell wall digestion can be skipped if it shows good FISH signal without this step.)
7. After cell wall digestion, enzymes were removed by three 2-min washes in DPBST.
8. Then samples were fixed in 4% PFA in PBS for 30 min at room temperature.
9. The fixative was removed by washing the samples in DPBST twice for 5 min.

### Sample permeabilization Monocots

1. Dehydrate the samples in a series of washes at RT with a tube revolver: 70% ethanol for 15 min, 90% ethanol for 15 min, 100% ethanol twice for 15 min each, 100% methanol twice for 15 min each. Leave samples in methanol at -20°C overnight. **PAUSE POINT: samples can be stored in Methanol at -20°C for several weeks before continuing.**
2. Incubate twice for 30 minutes in a solution of 100% Histo-Clear II at RT. Each time, apply vacuum for the first 10 minutes then transfer to a tube revolver for the last 20 minutes. Rehydrate the samples through a series of washes at RT with a tube revolver: 50% Histo-Clear II / 50% ethanol for 15 min, 100% ethanol for 15 min, 50% ethanol / 50% DPBST for 15 min - roots will float up then settle after a few minutes, then 100% DPBST twice for 15 min - roots will float up then settle after a few minutes.
3. Incubate with 4% formaldehyde in DPBST at RT under gentle vacuum in fume hood for 10 min. Fix the meristem for 20 min in 4% formaldehyde in DPBST at RT on a tube revolver.
4. Wash twice for 15 min each in DPBST at room temperature with a tube revolver.
5. Aliquot roots into 2 mL Eppendorf tubes. Use between 5-10 roots per tube/probe.

Proteinase K treatment can be performed after permeabilization if fluorescence from fluorescent proteins needs to be removed. Samples were treated with proteinase solution at 37 °C for 15 min and then washed by DPBST 3 times for 2 min. The digested samples were fixed in 4% PFA in PBS for 30 min and washed in DPBST twice for 5 min.

### HCR RNA-FISH Hybridization

HCR RNA-FISH was performed according to the previous publication (Choi et al., 2018) and online protocols provided by Molecular Instruments (https://www.molecularinstruments.com/hcr-rnafish-protocols).

Probe solution was prepared by adding 0.4 μL (1 μM stock) of each hybridization probe set into 100 μL pre-heated HCR Probe Hybridization Buffer (Molecular Instruments) at 37 °C.

#### Arabidopsis

Remove DPBST and replace with 200 μL pre-heated HCR Probe Hybridization Buffer (no probe). Samples were incubated in HCR Probe Hybridization Buffer for 30 min at 37 °C. Remove the Hybridization Buffer and add 100 μL probe solution. Samples were then incubated in probe solution overnight at 37 °C.

#### Monocots

Remove DPBST and replace with 500 μL of HCR Probe Hybridization Buffer (no probe). Apply gentle vacuum in fume hood for 10 minutes, then pre-hybridize by incubating for 1 hour at 37°C in a thermomixer with agitation.

## PAUSE POINT

### samples can be stored in HCR Probe Hybridization Buffer at –20°C for several weeks before continuing

Remove Hybridization Buffer and add the probe solution. Hybridize by incubating overnight (∼20h) at 37°C in a thermomixer with agitation.

Store an aliquot of HCR Probe Wash Buffer (Molecular Instruments) at 37°C overnight to pre-heat.

#### HCR RNA-FISH Amplification

After the HCR RNA-FISH hybridization, samples were washed with pre-heated HCR Probe Wash Buffer (Molecular Instruments) at 37 °C four times for 15 min. Then samples were washed by 5x SSCT twice for 5 min. For Arabidopsis, samples were then pre-amplified with 200 μL HCR Amplification Buffer (Molecular Instruments) at room temperature for 10 min. For Monocots, 5x SSCT was replaced by 500 μL HCR Amplification Buffer, and then gentle vacuum was applied in fume hood for 10 minutes. Monocots samples were pre-amplified in tube rotator at room temperature for 50 min.

While samples wash and pre-amplify, the hairpin solution is prepared. For Arabidopsis, 50 μL hairpin solution is prepared for each sample. For monocots, 250 μL hairpin solution is prepared for each sample. For 50 μL hairpin solution, 3 pmol hairpin h1 and 3 pmol hairpin h2 (i.e. 1 μL of the 3 μM stocks) were separately incubated at 95 °C for 90 seconds and cooled to room temperature in dark for 30 min. The hairpin solution is prepared by combining snap-cooled h1 and h2 hairpins in 50 μL of HCR Amplification Buffer at room temperature.

After the pre-amplification, the HCR Amplification Buffer was removed, and samples were incubated in hairpin solution overnight (∼20h) in the dark at room temperature.

Excessive hairpins were removed by washing in (1) 5x SSCT twice for 5 min, (2) 5x SSCT twice for 30 min, (3) 5x SSCT once for 5 min.

Samples can be kept in 5xSSCT for over one or two weeks at 4°C without any signal losses (depending on the probe brightness).

#### DAPI staining for Arabidopsis inflorescences

Samples were washed in DPBST once for 10 min before staining. To stain the nuclei, samples were incubated in DAPI staining solution for 10 min and washed by DPBS. Samples can be stored in DPBS at 4 °C in dark for at least several days before microscopy.

#### Sample mounting and microscopy

For Arabidopsis inflorescences, samples were mounted on 2% agarose gel in 60 mm petri dishes. The stem underneath the apex was gently inserted in agarose gel under a stereomicroscope using fine forceps. Mounted samples were submerged in water and imaged by an upright Leica Stellaris 5 White Light Laser confocal microscope equipped with a water immersion objective (HC PL APO 40×/1.10 W CORR CS2). A z-stack was captured from the top layer of the shoot apex to the deeper tissue. Details of confocal microscope settings for each figure were shown in Table 2. Maximum intensity projection, rotation, and orthogonal view were performed by FIJI (Schindelin et al., 2012). 3D projection in Fig. 2c is generated by Leica LAS X 3D Visualisation software.

Monocots: Transfer samples onto a glass slide (in 5x SSCT) and using a 15° microscalpel cut and arrange them so that the cut face of the roots is facing upwards before being covered with coverslip. Samples were imaged on Leica SPE inverted confocal microscope, using an air objective (20×/0.7) with 2.5× zoom within the LAS AF software (find details in Table 2).

**Table. 2.**
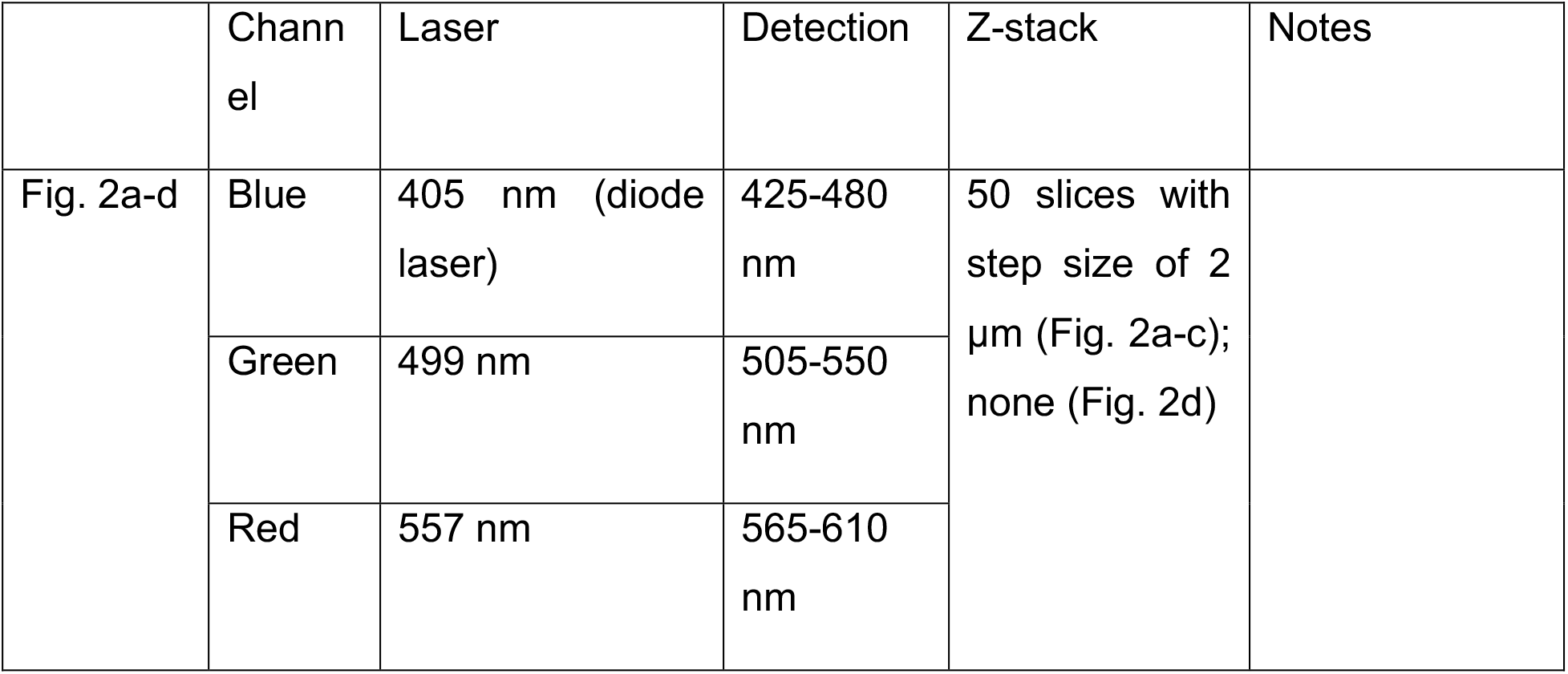
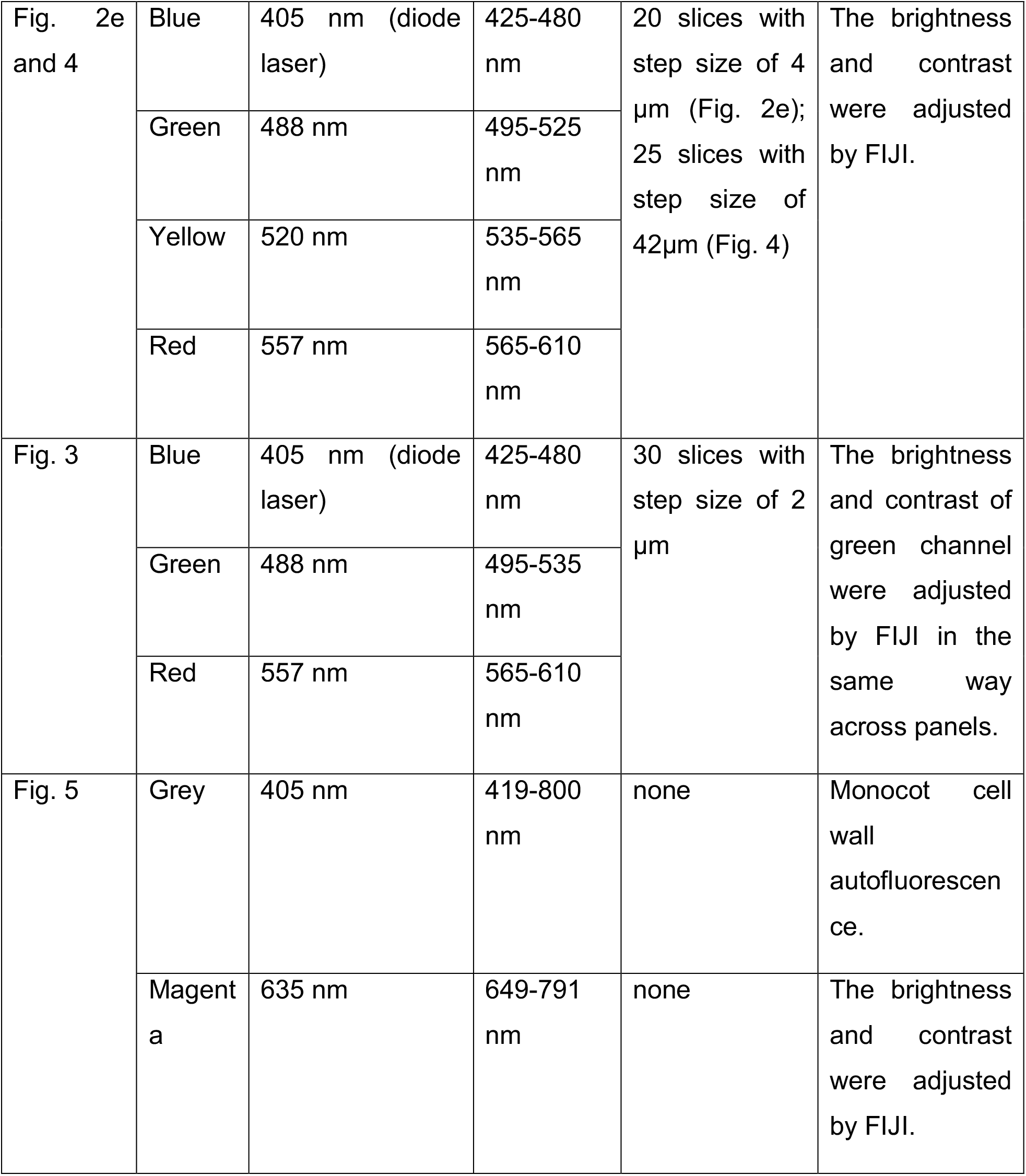
Confocal microscopy image acquisition settings

|  | Channel | Laser | Detection | Z-stack | Notes |
| --- | --- | --- | --- | --- | --- |
| Fig. 2a-d | Blue | 405 nm (diode laser) | 425-480 nm | 50 slices with step size of 2 µm (Fig. 2a-c); none (Fig. 2d) |  |
|  | Green | 499 nm | 505-550 nm |  |  |
|  | Red | 557 nm | 565-610 nm |  |  |
| Fig. 2e and 4 | Blue | 405 nm (diode laser) | 425-480 nm | 20 slices with step size of 4 μm (Fig. 2e); 25 slices with step size of 42μm (Fig. 4) | The brightness and contrast were adjusted by FIJI. |
|  | Green | 488 nm | 495-525 nm |  |  |
|  | Yellow | 520 nm | 535-565 nm |  |  |
|  | Red | 557 nm | 565-610 nm |  |  |
| Fig. 3 | Blue | 405 nm (diode laser) | 425-480 nm | 30 slices with step size of 2 μm | The brightness and contrast of green channel were adjusted by FIJI in the same way across panels. |
|  | Green | 488 nm | 495-535 nm |  |  |
|  | Red | 557 nm | 565-610 nm |  |  |
| Fig. 5 | Grey | 405 nm | 419-800 nm | none | Monocot cell wall autofluorescence. |
|  | Magenta | 635 nm | 649-791 nm | none | The brightness and contrast were adjusted by FIJI. |

#### Combined FISH and IHC

In combined FISH and IHC, HCR RNA-FISH was performed as indicated above, and IHC was immediately followed after HCR RNA-FISH. We adapted the IHC protocol from previous published IHC methods (Pasternak et al., 2015; Tran et al., 2021).

1. After HCR RNA-FISH, samples were fixed in 4% PFA in PBS for 15 min and washed by DPBST twice for 2 min.
2. To achieve better permeability for antibodies, the cell wall of the samples was further digested with cell wall digestion enzyme mix B for 5 min at room temperature.
3. Digested samples were fixed in 4% PFA in PBS for another 15 min and washed by DPBST twice for 2 min.
4. Samples were blocked in blocking buffer at 4 °C overnight with gentle rotation.
5. Then samples were incubated with primary antibody solution at 4 °C overnight with gentle rotation.
6. Excessive primary antibody was removed by washing in (1) DPBST once for 5 min, (2) DPBST twice for 30 min, (3) DPBST once for 5 min with gentle rotation.
7. Samples were incubated with secondary antibody solution at 4 °C overnight with gentle rotation.
8. Excessive secondary antibody was removed by washing in (1) DPBST once for 5 min, (2) DPBST twice for 30 min, (3) DPBST once for 5 min with gentle rotation.

DAPI staining and confocal microscopy for combined FISH and IHC were performed same as FISH samples as indicated.

## List of abbreviations

FISH: fluorescence in situ hybridization
HCR: hybridization chain reaction
DPBS: Dulbecco′s phosphate buffered saline
IHC: immunohistochemistry

## Acknowledgements

We thank Dr. Shalini Yadav for comments. Work on this method was supported by NSF IOS 1905062 to D.W. and Human Frontiers of Science (LT000972/2018-L) to B.G.

## Author Contributions

D.W. and K.D.B designed the research. T.H. conceived of the approaches and generated FISH and IHC in Arabidopsis thaliana, R.R and B.G developed and generated FISH in monocots.

## Reference

Barton, M.K. (2001). Giving meaning to movement. Cell 107, 129–132.

Birnbaum, K., Guillotin, B., Rahni, R., Passalacqua, M., Mohammed, M., Xu, X., Jackson, D., Groen, S., and Gillis, J. (2022). Gene duplication and cellular divergence in crops. Research Square. DOI: 10.21203/rs.3.rs-1739501/v1

Bradley, D., Ratcliffe, O., Vincent, C., Carpenter, R., and Coen, E. (1997). Inflorescence commitment and architecture in Arabidopsis. Science 275, 80–83.

Brewer, P.B., Heisler, M.G., Hejátko, J., Friml, J., and Benková, E. (2006). In situ hybridization for mRNA detection in Arabidopsis tissue sections. Nat. Protoc. 1, 1462–1467.

Choi, H.M.T., Beck, V.A., and Pierce, N.A. (2014). Next-Generation in Situ Hybridization Chain Reaction: Higher Gain, Lower Cost, Greater Durability. ACS Nano 8, 4284–4294.

Choi, H.M.T., Chang, J.Y., Trinh, L.A., Padilla, J.E., Fraser, S.E., and Pierce, N.A. (2010). Programmable in situ amplification for multiplexed imaging of mRNA expression. Nat. Biotechnol. 28, 1208–1212.

Choi, H.M.T., Schwarzkopf, M., Fornace, M.E., Acharya, A., Artavanis, G., Stegmaier, J., Cunha, A., and Pierce, N.A. (2018). Third-generation in situ hybridization chain reaction: multiplexed, quantitative, sensitive, versatile, robust. Development 145.

Choi, H.M.T., Calvert, C.R., Husain, N., Huss, D., Barsi, J.C., Deverman, B.E., Hunter, R.C., Kato, M., Lee, S.M., Abelin, A.C.T., Rosenthal, A.Z., Akbari, O.S., Li, Y., Hay, B.A., Sternberg, P.W., Patterson, P.H., Davidson, E.H., Mazmanian, S.K., Prober, D.A., van de Rijn, M., Leadbetter, J.R., Newman, D.K., Readhead, C., Bronner, M.E., Wold, B., Lansford, R., Sauka-Spengler, T., Fraser, S.E., and Pierce, N.A. (2016). Mapping a multiplexed zoo of mRNA expression. Development 143, 3632–3637.

Conti, L., and Bradley, D. (2007). TERMINAL FLOWER1 is a mobile signal controlling Arabidopsis architecture. Plant Cell 19, 767–778.

Dirks, R.M., and Pierce, N.A. (2004). Triggered amplification by hybridization chain reaction. Proc. Natl. Acad. Sci. U. S. A. 101, 15275–15278.

Duncan, S., Olsson, T.S.G., Hartley, M., Dean, C., and Rosa, S. (2016). A method for detecting single mRNA molecules in Arabidopsis thaliana. Plant Methods 12, 13.

Fletcher, J.C., Brand, U., Running, M.P., Simon, R., and Meyerowitz, E.M. (1999). Signaling of cell fate decisions by CLAVATA3 in Arabidopsis shoot meristems. Science 283, 1911–1914.

Goretti, D., Silvestre, M., Collani, S., Langenecker, T., Méndez, C., Madueño, F., and Schmid, M. (2020). TERMINAL FLOWER1 Functions as a Mobile Transcriptional Cofactor in the Shoot Apical Meristem. Plant Physiol. 182, 2081–2095.

Goto, K., Kyozuka, J., and Bowman, J.L. (2001). Turning floral organs into leaves, leaves into floral organs. Curr. Opin. Genet. Dev. 11, 449–456.

Gruel, J., Landrein, B., Tarr, P., Schuster, C., Refahi, Y., Sampathkumar, A., Hamant, O., Meyerowitz, E.M., and Jönsson, H. (2016). An epidermis-driven mechanism positions and scales stem cell niches in plants. Sci Adv 2, e1500989.

Hejátko, J., Blilou, I., Brewer, P.B., Friml, J., Scheres, B., and Benková, E. (2006). In situ hybridization technique for mRNA detection in whole mount Arabidopsis samples. Nat. Protoc. 1, 1939–1946.

Huang, K., Batish, M., Teng, C., Harkess, A., Meyers, B.C., and Caplan, J.L. (2020). Quantitative Fluorescence In Situ Hybridization Detection of Plant mRNAs with Single-Molecule Resolution. In RNA Tagging: Methods and Protocols, M. Heinlein, ed (New York, NY: Springer US), pp. 23–33.

Jack, T., Brockman, L.L., and Meyerowitz, E.M. (1992). The homeotic gene APETALA3 of Arabidopsis thaliana encodes a MADS box and is expressed in petals and stamens. Cell 68, 683–697.

Jonkman, J., Brown, C.M., Wright, G.D., Anderson, K.I., and North, A.J. (2020). Tutorial: guidance for quantitative confocal microscopy. Nat. Protoc. 15, 1585–1611.

Krizek, B.A., and Fletcher, J.C. (2005). Molecular mechanisms of flower development: an armchair guide. Nat. Rev. Genet. 6, 688–698.

Long, J., and Barton, M.K. (2000). Initiation of axillary and floral meristems in Arabidopsis. Dev. Biol. 218, 341–353.

Long, J.A., Moan, E.I., Medford, J.I., and Barton, M.K. (1996). A member of the KNOTTED class of homeodomain proteins encoded by the STM gene of Arabidopsis. Nature 379, 66–69.

Matsuyama, T., Satoh, H., Yamada, Y., and Hashimoto, T. (1999). A maize glycine-rich protein is synthesized in the lateral root cap and accumulates in the mucilage. Plant Physiol 120, 665–674.

Mayer, K.F., Schoof, H., Haecker, A., Lenhard, M., Jürgens, G., and Laux, T. (1998). Role of WUSCHEL in regulating stem cell fate in the Arabidopsis shoot meristem. Cell 95, 805–815.

Ortiz-Ramirez, C., Guillotin, B., Xu, X., Rahni, R., Zhang, S., Yan, Z., Coqueiro Dias Araujo, P., Demesa-Arevalo, E., Lee, L., Van Eck, J., Gingeras, T.R., Jackson, D., Gallagher, K.L., and Birnbaum, K.D. (2021). Ground tissue circuitry regulates organ complexity in maize and Setaria. Science 374, 1247–1252.

Pasternak, T., Tietz, O., Rapp, K., Begheldo, M., Nitschke, R., Ruperti, B., and Palme, K. (2015). Protocol: an improved and universal procedure for whole-mount immunolocalization in plants. Plant Methods 11, 50.

Rozier, F., Mirabet, V., Vernoux, T., and Das, P. (2014). Analysis of 3D gene expression patterns in plants using whole-mount RNA in situ hybridization. Nat. Protoc. 9, 2464–2475.

Schindelin, J., Arganda-Carreras, I., Frise, E., Kaynig, V., Longair, M., Pietzsch, T., Preibisch, S., Rueden, C., Saalfeld, S., Schmid, B., Tinevez, J.-Y., White, D.J., Hartenstein, V., Eliceiri, K., Tomancak, P., and Cardona, A. (2012). Fiji: an open-source platform for biological-image analysis. Nat. Methods 9, 676–682.

Seyfferth, C., Renema, J., Wendrich, J.R., Eekhout, T., Seurinck, R., Vandamme, N., Blob, B., Saeys, Y., Helariutta, Y., Birnbaum, K.D., and De Rybel, B. (2021). Advances and Opportunities in Single-Cell Transcriptomics for Plant Research. Annu. Rev. Plant Biol. 72, 847–866.

Shaw, R., Tian, X., and Xu, J. (2021). Single-Cell Transcriptome Analysis in Plants: Advances and Challenges. Mol. Plant 14, 115–126.

Smyth, D.R., Bowman, J.L., and Meyerowitz, E.M. (1990). Early Flower Development in Arabídopsis. Plant Cell 2, 755–767.

Solanki, S., Ameen, G., Zhao, J., Flaten, J., Borowicz, P., and Brueggeman, R.S. (2020). Visualization of spatial gene expression in plants by modified RNAscope fluorescent in situ hybridization. Plant Methods 16, 71.

Tran, T.M., Demesa-Arevalo, E., Kitagawa, M., Garcia-Aguilar, M., Grimanelli, D., and Jackson, D. (2021). An Optimized Whole-Mount Immunofluorescence Method for Shoot Apices. Curr Protoc 1, e101.

Trivedi, V., Choi, H.M.T., Fraser, S.E., and Pierce, N.A. (2018). Multidimensional quantitative analysis of mRNA expression within intact vertebrate embryos. Development 145.

Yang, W., Schuster, C., Prunet, N., Dong, Q., Landrein, B., Wightman, R., and Meyerowitz, E.M. (2020). Visualization of Protein Coding, Long Noncoding, and Nuclear RNAs by Fluorescence in Situ Hybridization in Sections of Shoot Apical Meristems and Developing Flowers. Plant Physiol. 182, 147–158.

Yanofsky, M.F., Ma, H., Bowman, J.L., Drews, G.N., Feldmann, K.A., and Meyerowitz, E.M. (1990). The protein encoded by the Arabidopsis homeotic gene agamous resembles transcription factors. Nature 346, 35–39.

Young, A.P., Jackson, D.J., and Wyeth, R.C. (2020). A technical review and guide to RNA fluorescence in situ hybridization. PeerJ 8, e8806.

Zhu, Y., Klasfeld, S., Jeong, C.W., Jin, R., Goto, K., Yamaguchi, N., and Wagner, D. (2020). TERMINAL FLOWER 1-FD complex target genes and competition with FLOWERING LOCUS T. Nat. Commun. 11, 5118.

